# The domesticated brain: genetics of brain mass and brain structure in an avian species

**DOI:** 10.1101/066977

**Authors:** R. Henriksen, M. Johnsson, L. Andersson, P. Jensen, D. Wright

## Abstract

As brain size usually increases with body size it has been assumed that the two are tightly constrained and evolutionary studies have therefore often been based on relative brain size (i.e. brain size proportional to body size) instead of absolute brain size. The process of domestication offers an excellent opportunity to disentangle the linkage between body and brain mass due to the extreme selection for increased body mass that has occurred. By breeding an intercross between domestic chicken and their wild progenitor, we address this relationship by simultaneously mapping the genes that control inter-population variation in brain mass and body mass. Loci controlling variation in brain mass and body mass have separate genetic architectures and are therefore not directly constrained. Genetic mapping of brain regions in the intercross indicates that domestication has led to a larger body mass and to a lesser extent a larger absolute brain mass in chickens, mainly due to enlargement of the cerebellum. Domestication has traditionally been linked to brain mass regression, based on measurements of relative brain mass, which confounds the large body mass augmentation due to domestication. Our results refute this concept in chicken and confirm recent studies that show that different genetic architectures underlie these traits.

## INTRODUCTION

Brain size variation across vertebrate species continues to fascinate evolutionary biologists, due to the cognitive and behavioral phenotypes it is thought to underlie. Most studies on brain size differences suggest some kind of trade-off between the costs of developing and maintaining energetically expensive brains and certain physiological variables (such as body size^1^, metabolic rate^2^, development time^3^) or lifestyle variables (e.g. foraging ecology^4^ and social environment^5^). One physiological variable that correlates notably with brain size is body size^1^. As brain size usually increases with increasing body size^6^ it has been assumed that the two are tightly constrained during developmental growth^7^. Researchers have therefore often relied on relative rather than absolute brain size in correlative studies. The relationship between body size and brain size is, however, poorly understood and the use of allometry in brain size evolution studies has been criticized^8–10^. Understanding the genetics of brain size evolution is extremely pertinent to determine the relationship between brain size and body size. Most importantly, to what degree there is overlap (and potential pleiotropy) between the genes responsible for both. To date, studies on the genetic relationship between brain and body size are almost entirely limited to phylogenetic comparisons and measures of selection, and have failed to identify the overlap of the genetic architecture between these traits (brain size and body size), especially using a within-species approach. The analysis of the rates of evolution in a between-species analysis of cichlids can indicate that brain size and body size can have distinct rates of evolution^11^. Similarly, selection on body mass can potentially drive reduced relative brain mass^12^, whilst different orders of animals may have different brain-body mass variation, which is driven primarily by variability in body mass^13^. Genetic correlations between brain and body size in stickleback (*Gasterosteus aculeatus*) showed a positive correlation between the two traits^14^, but also a large standard error to this estimate (with the correlation being between 12-96%). In the principle study to actually assess the genetic architecture of quantitative variation in brain size, a quantitative trait loci (QTL) study in inbred mice strains identified genomic regions associated with overall brain mass^15^, though was primarily focussed on identifying QTL for separate brain sub-regions. In this study, however, the genetic architecture for body mass was almost entirely unresolved (identifying only one suggestive locus), making it impossible to assess any potential overlap between brain and body mass loci. Therefore, nothing is known regarding the combined genetic architectures for brain mass and body mass. Thus, to date, no studies exist that comprehensively examine the genetic architecture of both brain mass and body mass in an intraspecies specific manner.

The genetics underlying loci affecting overall brain size and body size in avian (or indeed any other) species have yet to be explored, although the genes underpinning certain brain regions in birds have been investigated. Most genetic work on avian brain regions has been on genes relating to variation in the brain structures governing song learning^16–18^, and genetic programs involved in determining the basic architecture of the telencephalon^19–21^. The genetic basis of overall brain composition differences has yet to be investigated in birds. The degree to which different brain regions can develop independently is highly debated. According to the ‘mosaic evolution’ hypothesis individual brain regions can develop and grow independently in size^22^ while the ‘concerted evolution’ hypothesis argues that different brain regions have been limited by developmental constraints and that brain size alters predominantly as a whole^3^. In the case of the latter hypothesis therefore individual regions cannot change in size independently. To date, only two studies have attempted to identify gene regions that affect variation in brain substructure size within species. The previously mentioned QTL study in inbred mice strains^15^, and a Genome Wide Association Study (GWAS) study by Hibar et al.^23^ that identified eight loci affecting putamen and caudate nucleus volumes in humans.

Brain size differs substantially within species^24^. Since evolution operates through intraspecific variation, within species differences can help disentangle the relationship between brain and body size and the degree to which different brain structures are constrained developmentally. The process of domestication is especially interesting because of the huge differences in both brain and particularly body size in domesticated animals as compared to their wild progenitors, and the reduction in relative brain size. This generates a perfect model for assessing the effects of variation in brain and body size, how body size is constrained by brain size and vice-versa. A classical effect of domestication is reduced relative brain size, with this believed to reflect the reduced functional needs of domesticated animal’s brains. Studies have reported smaller brain size in both domesticated mammalian (sheep:^25^, pig:^26,27^, mink:^28^), and avian species (turkeys:^29^, chickens:^30^, pigeons:^31^, ducks:^32^), as compared to their wild progenitors. However, the methods used in virtually all studies on brain size reduction in domesticated animals confound the effect of brain reduction with the often very large body size augmentation that has occurred during domestication. The effect of reduced relative brain size with directional selection is not limited to domestication, with studies on fish showing that reduced relative brain mass can be potentially driven by increased selection for body mass^12^. Thus results from the genetic dissection of domestication may be pertinent in a wider perspective.

Genomic sequencing of the wild progenitor of all domesticated chickens, the Red Junglefowl (RJF) *Gallus gallus*, as well as different domestic chicken breeds has enabled the identification of selective sweeps – gene regions that have been strongly selected upon during domestication^33^. Although the domestic chicken and RJF differ both physiologically and behaviourally they still belong to the same species and can interbreed. This allows us to perform a similar QTL analysis as that by Hager and colleagues in mice^15^ but with the added advantage that large differences in the domestic chicken and RJF brain and body mass and genome provides us with an *a priori* hypotheses regarding brain and body phenotype (mass) and which genes might be involved (underlying selective sweep-regions).

In this study we use a domestic (White Leghorn chicken, *Gallus gallus domesticus*) x wild (RJF) advanced intercross to fine-map quantitative trait loci (QTL) pertaining to brain mass and body mass differences between wild and domestic populations, as well as ontogenetical analyses of brain and body mass development in wild and domestic birds. This intercross allows us to separately map both brain mass and body mass QTL and determine the genetic architecture of both traits. Brains from this intercross population were further subdivided into 4 brain regions and weighed. To access potential differences in cerebrotype, each brain region was mapped both as a whole-weight measure and in terms of the proportion of total brain mass accounted for by each region (termed ‘proportional QTL’). It is important to note that the QTL mapping procedure only detects loci that differ between the parental populations, i.e. those loci that are distinct between wild and domestic birds. Loci that both populations have in common will not be detected. In this way, we can therefore specifically detect those loci that control brain mass and body mass that have been selected upon by domestication. However, the general loci that both domestic and Red Junglefowl share will be undetected, and therefore missed. Ontogenetic brain and body mass development analyses were performed using populations of wild and domestic birds. These allow the developmental growth trajectories of both brain mass and body mass to be compared between wild and domestic birds. If brain mass is indeed inextricably linked with body mass, the growth trajectories of both brain mass and body mass should mirror one another, in both wild and domestic populations. This enables a separate analysis of the relationship between brain and body mass in an inter-population context. To our knowledge, this is the first study to actually identify multiple genomic regions underpinning the mass differences in both whole brain and brain region mass, whilst simultaneously mapping body mass, not only between a wild and domesticated population but also in vertebrates in general. This approach separates and decouples the detected brain loci at a genetic level from those loci affecting only body mass.

## METHODS

### Chicken Study population and cross design

The intercross population was an eighth generation intercross between a population of RJF, derived originally from Thailand^34,35^ and a line of selected White Leghorn (WL) chickens, with a total of 470 adult F_8_ individuals were used in this study. Females were assayed for brooding prior to culling. In addition a total of 61 RJF and 65 WL chickens were reared (in conditions identical to the intercross birds) to assess brain development and growth differences from age one, two, four, ten and fifteen weeks as well as at adult age, between RJF and WL. Eleven broiler birds were additionally dissected at two weeks of age, as a comparison for the other major domesticated strain. Volumetric measurements were taken on the brains of two adult male RJF and 2 adult male WL individuals. The study was approved by the local Ethical Committee of the Swedish National Board for Laboratory Animals. All methods were performed in accordance with the relevant guidelines and regulations.

### Phenotyping

#### Brain measurements and dissection

Immediately after culling, the brains were removed from the birds and weighed, after being dissected into four pieces for 315 individuals and nine pieces for 129 individuals. Total brain mass values were available for 439 of these individuals. Four piece dissections involved dividing the brain into the cerebral hemispheres, optic tectum, cerebellum and a brainstem region (which included thalamus, the rest of the midbrain and the hindbrain; for more information on the dissected brain regions see supplementary figure 1 and the supplementary methods). Nine-piece dissections were only used for QTL analysis of body mass and whole brain mass, with these dissections being used for a different study (^36^ – with the brainstem region further subdivided into the optic chiasma, thalamus, hypothalamus, medulla oblongata, pons and nucleus tractus solitarii). Each brain region was weighed to the nearest 0.001g. To ascertain whether a heavier brain also equates to a larger brain, brain mass was compared with volume in a subset of RJF and WL birds. The proportional brain region mass was calculated by dividing the mass for each specific region by the total brain mass. Details of volumetric measures are given in the supplementary methods section as is the technique for assessment of brooding. Brain volume was found to scale linearly with mass (see supplementary figure 2A). In all four brain regions, regions with a larger mass had a greater volume, with brain mass explaining 98-100% of the variation when regressed on brain volume (supplementary figure 2B).

#### Genotyping, QTL and mapping

All individuals were genotyped for 652 SNP markers, with QTL analysis performed using R/Qtl^37^ for both standard interval mapping and epistatic analyses. See supplementary methods for full details of the mapping and models used.

## RESULTS

### Total brain mass is larger in domestic birds with a different growth trajectory relative to body mass in comparison to wild birds

Absolute brain mass is larger in domestic (WL) chickens relative to wild RJF (figure 1), with a consistent difference of approximate ~0.2g up to 5 weeks of age, with this increasing to ~0.4g at sexual maturity. In the case of both domestic chickens and RJF brain growth is linear up to adulthood. In contrast body mass differences diverge rapidly after 4 weeks of age between domestic chickens and RJF, with domestic chickens showing a sharp exponential increase in mass, and at sexual maturity they weigh almost twice as much as RJF (figure 1). Thus whilst domestic chickens exhibit the classic effects of domestication (reduced relative brain mass, massively increased body mass), the differences between brain growth and body growth trajectories between these two populations suggests that different genetic systems are at play governing these different traits. If this is the case, then the genetic architecture of brain mass and body mass in the intercross population should be non-overlapping. Additionally, the brain composition of wild birds and domestic birds was also found to differ, with domestic birds possessing a larger cerebellum and cerebral hemispheres and smaller optic tecta and brainstem with these differences being largely consistent throughout post-hatch growth (Supplementary figure 3). Therefore, non-overlapping architecture for different brain sub-regions in the intercross would also favour the mosaic brain theory over the concerted brain hypothesis.

**Figure 1.**
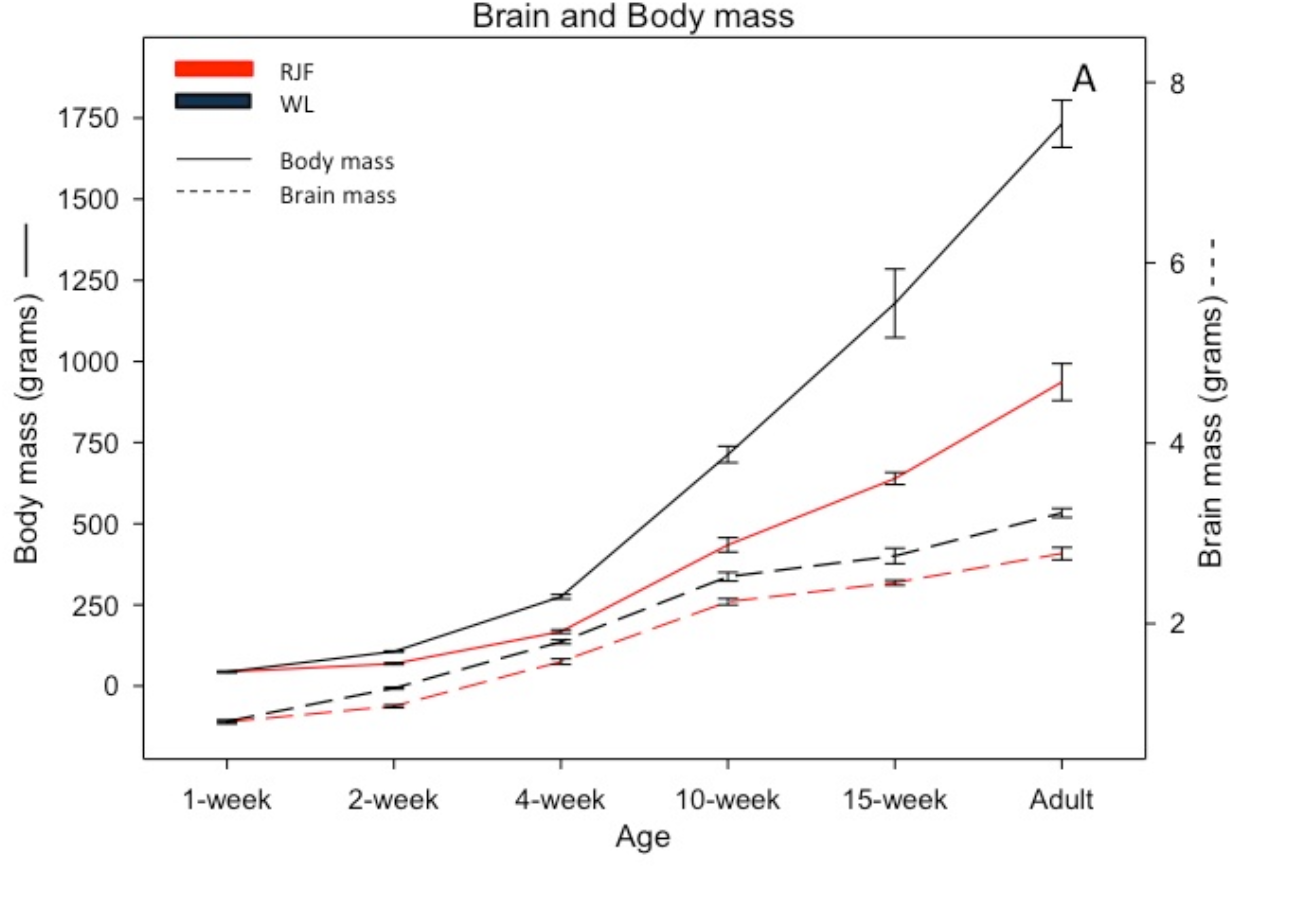
Changes in body mass (solid lines: mean±SE) and brain mass (dotted lines: mean±SE) in White leghorn (black lines) and Red Junglefowls (red lines) from 1-week of age until adulthood.

### The detected architectures for inter-population brain mass and body mass appear separate

A total of 20 QTL relating to inter-population brain mass and proportional brain mass variation were identified (table 1 and supplementary table 2). One of the first things of note is that the QTL for brain mass and proportional brain region mass and absolute brain region mass are entirely separate from the QTL for body mass. There is no overlap for any of the loci involved, indicating that the genes selected by domestication (i.e. those leading to inter-population variation) for body mass and brain region mass are at least partially separate, i.e. it is possible to select for increased brain growth without increased body mass and vice-versa during domestication (see figures 2 and 3). Although body mass was not used as a covariate for the brain mass QTL analysis presented above (due to the possibility that including it could mask QTL that overlap body mass by already factoring body mass out the model), its inclusion lead to no changes in the detected QTL position, and in fact strengthened the QTL in some cases, further highlighting the separate nature of the two architectures. A general caveat with QTL analysis is that the genes controlling brain mass and body mass that are shared between domestic and wild populations (i.e. intra-population variation) will not be detected as QTL in this analysis. However, the phenotypic correlation between brain and body mass in this cross is relatively low and undetected loci causing intra-population variation that also overlapped would increase this correlation. Although significant, sex and body mass together account for only 18% of the total variation in brain mass, with sex alone accounting for 16% of the variation. Similarly, when a linear model fitting all the detected QTL as well as body mass was regressed onto brain mass, body mass still only accounted for 2% of the variation in brain mass variation, whilst the detected genetic loci accounted for 22% of the brain mass variation. A further caveat is that genes with effects too small to be detected may yet be pleiotropic between brain and body mass, though given the lack of any phenotypic correlations as noted above, this seems less probable. In the case of the genetic architecture for body mass, the detected QTL account for over 36% of the total variation for body mass (excluding sex differences), whilst the total GLM that included all body mass QTL and other covariates explained 84% of the variation for body mass. Therefore, relatively little unexplained variation is present in the model for body mass. A significant sex interaction can indicate that a QTL has greater effect on one sex as compared to the other. In this case, very few sex interaction effects were seen for cerebellum and total brain mass traits, with the only interactions being on chromosomes 3 (total brain mass) and 5 (proportional cerebellum mass), and none were found for any other traits (see supplementary table 3).

**Figure 2.**
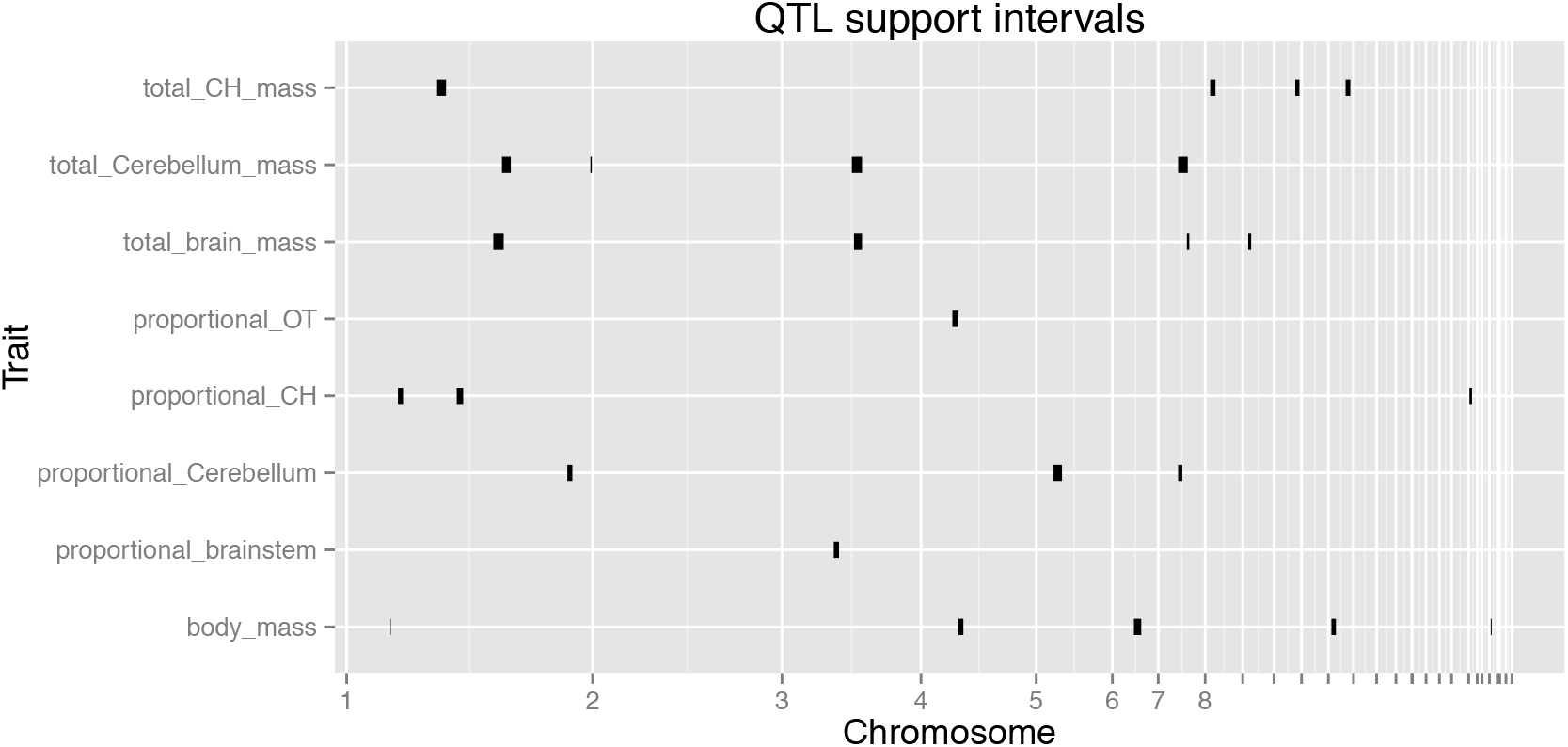
QTL supports intervals (as determined by the 1.8 LOD drop method) in the genome separating out loci for proportional and total brain mass and body mass. X-axis represents the chromosome containing the QTL, whilst the size of the each bar represents the total QTL region size (i.e. the 95% C.I. of the QTL location).

**Figure 3.**
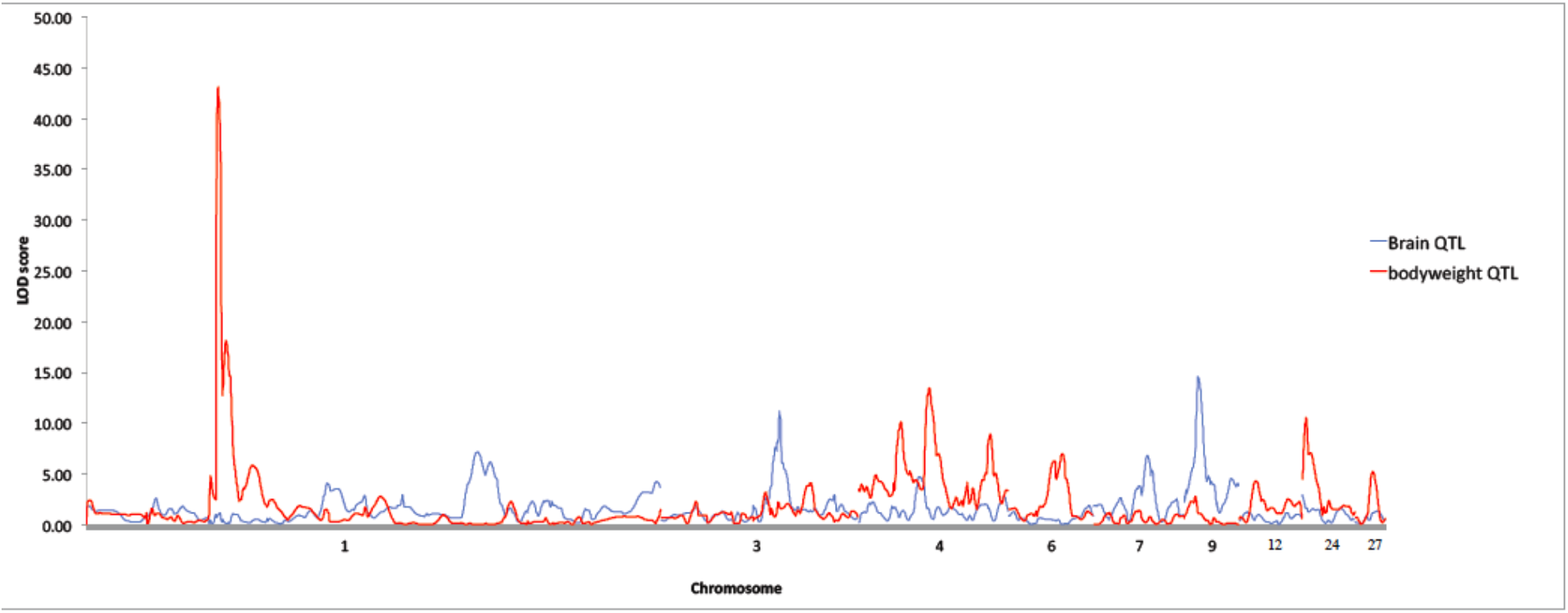
Comparison of LOD graphs for total brain mass QTL and body mass QTL

**Table 1.**
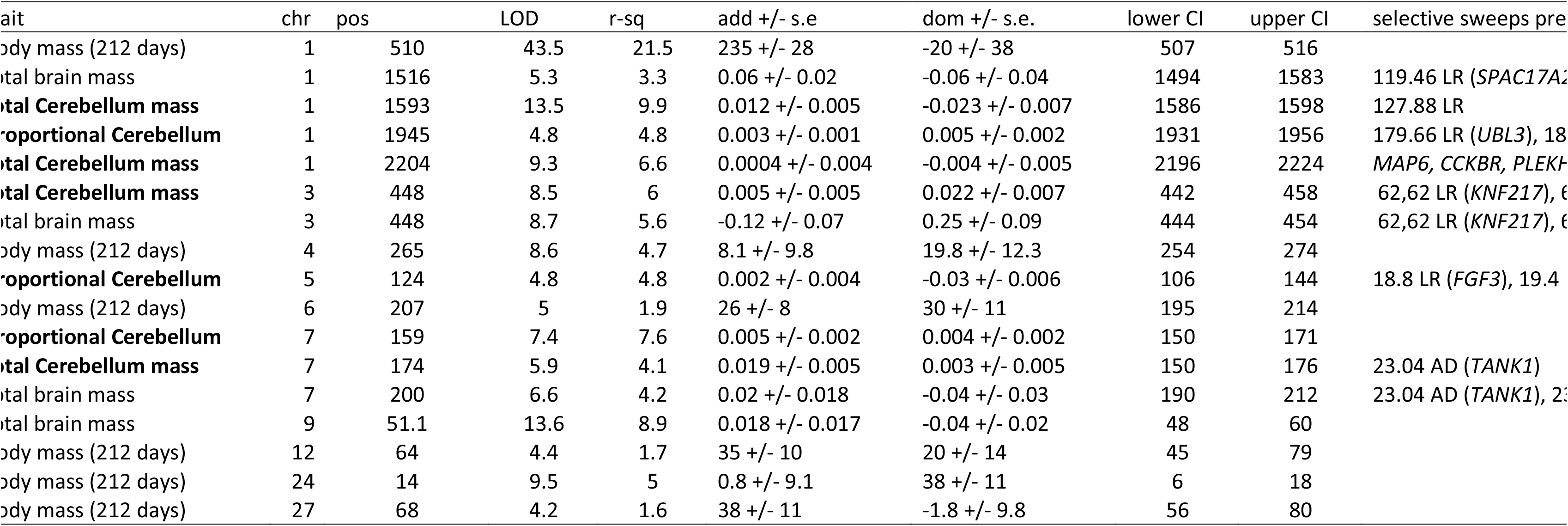
QTL information for body mass, total brain mass and cerebellum QTL. Includes locations (both the chromosome and the position in centiMorgans), % variance explained by each QTL (r-squared), additive and dominance effect sizes (positive values for additive values indicate a larger QTL effect size in domestic genotype birds, negative a larger value in wild genotype birds). The lower and upper bounds of the 95% confidence interval (C.I) are noted. The total QTL region is therefore the region bounded between these two limits. Locations of selective sweeps are also provided, with AD indicating the sweep is present in both Broiler and Layer birds, and LR indicating the sweep is specific to Layer birds. For sweeps present in cerebellum and total brain mass QTL any genes present within sweeps are also provided after the sweep location. Cerebellum QTL are marked in bold.

### Domestic genotypes cause larger brains and larger cerebella than wild genotypes

In the case of the cerebellum and total brain mass QTL, the observed allelic effect is in accordance with the phenotypic differences seen in the wild and domestic breeds (see supplementary figures 3-4). Domestic alleles correlate with an increase in cerebellum mass and total brain mass in 6 out of 7 of the cerebellum QTL and 3 out of 4 of the brain mass QTL identified, i.e. the QTL are not transgressive (see the additive and dominance QTL effect sizes in table 1), meaning that the alleles correlating with an increase in the phenotype come from the parental strain possessing the larger phenotype. In combination with the observation that brain mass has a separate genetic architecture to body mass in this cross, this indicates that once the genes underpinning brain mass and body mass are de-coupled, domestic chickens have larger, not smaller, brains as compared to their wild counterparts. Further to this, the cerebellum is also larger in mass in the domestic chickens (both totally and as a proportion of total brain mass), with the genetic control of this trait separate from overall body mass. There is a strong overlap between the observed genetic architectures for the cerebellum and total brain mass, with three of the four brain mass QTL overlapping total cerebellar mass QTL (on chromosomes 1,3, and 7, see table 1).

### Brain regions each have their own unique genetic architecture

The different brain regions have separate genetic architectures from one another for both total and proportional mass, as determined by the loci detected. Although in the case of the optic tectum and brainstem regions very few QTL were identified (only two loci in total). For both cerebral hemispheres and the cerebellum multiple QTL were identified (18 in total), explaining a relatively large amount of the variation in the cross (26% of total variation for proportional cerebral hemispheres mass, 26% of total variation for total cerebellum mass, 22% of the variation for total brain mass). Even with these more complete genetic architectures, there was nevertheless no overlap between different brain regions QTL with one another, indicating that, at least partially, this supports the mosaic theory of brain evolution^22^.

### Brain mass is selected for during domestication

Selective sweeps have been putatively identified in the chicken genome, representing regions that have reduced heterozygosity during the periods of selection that occurred during domestication^33^. By using a clustering analysis based on the number of sweeps detected in the genome, the confidence intervals of the brain QTL and the number of overlaps between brain QTL and selective sweep loci, we found that the QTL relating to cerebellum mass (7 loci in total, marked in bold in table 1) contained a significant enrichment of selective sweeps (P=0.04, using the clustering test as described in the supplementary methods), as did the regions relating to total brain mass (4 loci overlapped, P=0.01, clustering test). These results indicate that loci affecting cerebellum and total brain mass may have been directly selected upon during domestication, or that these loci are closely linked with other genes targeted by domestication selection. Given this significant overlap between selective sweeps and QTL regions, genes contained within these sweeps are good candidates for the genes causal to the QTL (often referred to as Quantitative Trait Genes). Twelve genes are present in the selective sweeps within total brain and cerebellar QTL (see table 1). Four of these (*FGF3, FGF9, BIN1, SHANK1*) have previous associations with neuronal conditions or neurogenesis (see discussion).

### Brooding behaviour correlates with proportional cerebellum mass

Given the strong selection on the mass of the cerebellum in domestic chickens, we correlated brooding behaviour with proportional cerebellum mass. We found a strong negative relationship between brooding behaviour and proportional cerebellum mass (GLM, n=123, F=−2.6, P=0.009), indicating that birds with a larger cerebellum and thereby a more domesticated brain cerebrotype displayed less brooding behaviour, characteristic of domestic chickens. Body mass was included in the model as a covariate, so these results are not due to overall body mass effects. Four QTL were detected for brooding behaviour (see supplementary table 2). Although none overlapped total brain mass QTL, the confidence interval of one (on chromosome 9) was within 32cM of a total brain mass QTL, whilst another (on chromosome 4) was within 38cM of a QTL for proportional optic tectum mass. In the case of the latter this distance corresponds to around 200kb, whilst in the case of the latter it is around 10Mb. No correlations were found between other brain regions and brooding.

### RJF and WL fat/ lean/ bone body tissue proportions

The body composition, and in particular the lean mass to fat mass percentage, could potentially be important in explaining intra-species brain mass differences, especially if lean mass requires a proportional increase in neurons. If domestic birds have a far greater percentage of fat mass, then departures from the usual brain allometry may be due to large brain mass not being required for the extra fat tissue. Using data from a previous study by Rubin and colleagues^38^ that measured overall fat, lean and bone mass in both wild and domestic chickens using Dual X-ray Absorbancy (DXA) techniques it is possible to calculate the relative proportions of these tissue types (see supplementary table 4). Although domestic females have a higher percentage of body fat (65% lean mass in WL females to 84% lean mass in RJF females), the domestic males (that show the largest decrease in relative brain mass) actually possess a far lower percentage body fat than their wild counterparts, with a correspondingly higher proportion of lean muscle mass (86% lean mass in WL males to 77% lean mass in RJF males).

## DISCUSSION

Domesticated chickens (WL) have a larger brain mass and body mass than their wild progenitor, but whereas body mass has increased by ~85% during domestication, brain mass has only increased by ~15%. This indicates that brain mass has been altered less by selection during domestication than body mass and that in chickens reduced relative brain mass during domestication has mainly been caused by an increase in body mass. By separating the linkage between loci affecting inter-population variation in brain mass and body mass in an advanced intercross we demonstrate that domestication selection has acted on apparently separate loci to increase brain mass and body mass in domesticated individuals. Therefore selection on body mass is not limited by brain mass and vice-versa in the chicken, with domestication leading to (at least partially) separate genetic architectures for these traits. It is important to note that brain and body mass architectures that are common to both wild and domestic populations will not be detected with this analysis. This means we cannot infer that no loci affecting brain mass and body mass are pleiotropic or overlap, only that those that are due to domestication selection do not. Similarly, with any QTL study there is always the possibility that loci with effect sizes too small to detect exist and do overlap and exhibit pleiotropy. However, the relatively small percentage of variation of brain mass explained by body mass suggests that these within-population loci are of smaller effect than the between-population loci. In the combined model of QTL and body mass covariates affecting brain mass, body mass explains only 2% of brain mass variation whilst the QTL explains around 22%, and sex explains 4% of the variation. Likewise, if numerous undetected, small-effect loci were also pleiotropic we would still expect the brain and body mass correlation to be higher. It is also possible that developmental constraints between brain and body mass may not necessarily exist at a pleiotropic level, but there could still be a physiological constraint. In this case, however, a stronger correlation between brain mass and body mass would still be expected in the above linear model. Our study is the first to provide evidence for a relatively distinct genetic architecture for body mass and brain mass in an avian species and also to demonstrate that separate loci underlie the mass of the different brain regions, thereby showing that the mass of an individual brain regions can be selected upon without being strictly constrained by the mass of the other brain regions, as predicted by the mosaic brain evolution hypothesis. Support for mosaic brain evolution has also been reported in mammals^15^ and fish^14^ and together with our findings in birds suggest that mosaic brain evolution is possible across vertebrate species.

The nature of the body and brain mass increases during domestication is of relevance when considering allometric scaling and the relationship between brain mass and body mass. It has been proposed that the developmental constraints limiting brain and body mass are related to the overall musculature (see the trophic theory of neural connections^39^, amongst others), with increased neural circuitry required for an increase in muscular anatomy. A possibility therefore is that domestic birds have a lower relative brain mass due to increased fat reserves making up a larger percentage of their overall body mass. The majority of the mass gain in domestics would then be due to increased fat deposits and thus not require any increase in brain mass. However, the calculations we performed using the data from a previous study by Rubin and colleagues^38^ indicates that this does not appear to be a limiting factor in this instance, with male WL birds having a higher lean body mass percentage than their RJF counterparts. Similarly, we have also shown that in chickens an increase in brain mass also correlates with an increase in brain volume. Similar strong correlations between brain mass and brain volume have recently been shown in stickleback fish^14^. This suggests that the density of brain tissue is constant, however it remains an open and intriguing question whether the increased brain mass induced by domestication relates to the incorporation of more neurons, or alternatively non-neuronal cells such as blood cells or glia, connections and the like.

Although the brain and the four different brain regions grow continuously until adulthood in both RJF and WL, the proportional mass of each brain region changes during posthatch development and domestic and wild birds differ in the proportional mass of certain brain regions. These differences in cerebrotype between RJF and WL, caused by selection during domestication, must therefore occur initially during prehatch development or during the first week posthatch. Our findings suggest that selection during domestication in chickens has been stronger on the cerebral hemispheres and cerebellum than on the brainstem region and optic tectum. This is further supported by the enrichment of selective sweeps (gene regions that have been strongly selected upon during chicken domestication), which are present in the QTL regions for both cerebellum and overall brain mass. Increased proportional cerebellum mass, compared to their wild counterparts, is seen in most studies on domesticated birds, including domesticated geese^40^, turkeys^29^ and pigeons^41^ (but see also^31^ for opposite results), but not in ducks^32^, suggesting that proportional enlargement of the cerebellum may have played an important role during domestication in several birds species.

We find that brooding behaviour is inversely correlated with the proportional mass of the cerebellum. These findings suggest that the cerebellum could help govern this behaviour, and therefore provide a link between domestication effects on brain composition (an enlarged cerebellum) and domestication effects on behaviour (reduced broodiness). Proportional enlargement of the cerebellum and cerebral hemispheres could potentially have been important for the chicken to adapt to several aspects of the domesticated environment. In birds increased cerebral hemispheres mass has been linked with increased social complexity^42^ and the cerebellum has been linked with foraging strategy^43^. Wild RJF live in small group sizes of around 4-10 individuals^44^, whereas domestic chickens are kept in far larger groups. Although these theories are tempting to extrapolate upon, care must be taken in their interpretation given the purely correlative data.

The ultimate goal of a QTL study is often to identify the genes themselves that underlie the trait in question. In standard (F_2_ or similar) QTL analyses, the confidence intervals of detected QTL are so large (typically in the region of 20-30 megabases) that identifying putative candidate genes is virtually impossible. However, in the study presented here the combination of the advanced intercross and the overlap with selective sweeps for domestication yield a number of candidate genes for brain growth. In regards to the QTL for cerebellum mass and total brain mass, 12 genes in total are present within sweeps in these regions. Of these, four have prior associations with neural conditions and neuronal generation, reinforcing them as excellent candidate genes for increased brain and cerebellum mass. The chromosome 1 sweep at 182.6Mb, in the middle of the QTL region for total cerebellum mass, contains the gene *FGF9*, which regulates the generation and positioning of Bergmann glia cells in the developing mouse cerebella^45^. *FGF9* also has a crucial role in embryonic neurological development^46–49^. Similarly the QTL for proportional cerebellum mass on chromosome 5 contains 4 sweeps, with three genes present in those sweeps. Once again an *FGF* gene, in this case *FGF3*, is present. Also present in a separate sweep is *SHANK1*, mutations in which are associated with dysfunction of glutamatergic synapses that lead to a variety of neuropsychiatric disorders including autism and schizophrenia^50^. The sweep located at 25.42Mb on chromosome 7 that overlaps with the total brain mass QTL is located within 100kb of the gene *BIN1* that has effects on working memory, hippocampal volume and functional connectivity^51^. The QTL for total cerebellum mass on chromosome 1 at 2204cM has an interval of 1.1Mb and is therefore sufficiently highly resolved to also address the genes contained within for potential functionality, containing as it does 15 reference sequence genes. Of these, a number of highly interesting genes are identified for further investigation. *MAP6* is in the centre of the QTL confidence interval and mediates neuronal connectivity for axonal growth^52^. It has been linked with synaptic plasticity anomalies and has associations with schizophrenia through neuronal transport defects^53^ and cognitive impairment^54^ *CCKBR* is linked with posttraumatic stress disorder and synaptic plasticity^55^ and *PLEKHB1* is associated with Attention Deficit Hyperactivity Disorder^56^.

Our findings reinforce the concerns from recent years regarding the use of relative brain mass in evolutionary studies. Although brain mass can co-vary with body mass, and allometric effects still exist, if the differing genetic architectures are confounded together meaningful differences will be masked. Most notably, selection can increase body mass irrespective of brain mass, and vice versa, so the use of relative measures can potentially be flawed. Our results also refute the common notion that domestication leads to the regression of brain mass in domesticated chicken as a result of reduced functional needs. The combination of an advanced intercross and the presence of selective sweeps give a number of high confidence candidate genes with putative effects on total brain and cerebella growth. This demonstrates that the RJF and the domestic chicken provide an interesting animal model for studying brain mass evolution during domestication, and also a general model for studying evolution in brain mass and composition.

## AUTHORS’ CONTRIBUTIONS

Conceived and designed the experiments: DW PJ LA RH. Performed the experiments: DW RH. Analyzed the data: RH MJ DW. Contributed reagents/materials/analysis tools: DW LA PJ. Wrote the paper: RH MJ DW.

## COMPETING FINANCIAL INTERESTS

All authors have no competing interests.

## DATA ACCESSIBILITY STATEMENT

Full genotype and phenotype data is available on dryad with the following doi: XXXXX.

## ACKNOWLEDGEMENTS

The research was carried out within the framework of the Swedish Centre of Excellence in Animal Welfare Science, and the Linköping University Neuro-network. SNP genotyping was performed by the Uppsala Sequencing Center. The project was supported by grants from the Carl Tryggers Stiftelse, Swedish Research Council (VR), the Swedish Research Council for Environment, Agricultural Sciences and Spatial Planning (FORMAS), and European Research Council (advanced research grant GENEWELL 322206).

